# 300 Hz transcutaneous auricular vagus nerve stimulation (taVNS) impacts pupil size nonlinearly as a function of intensity

**DOI:** 10.1101/2024.04.19.590334

**Authors:** Ian Phillips, Michael A. Johns, Nick B. Pandža, Regina C. Calloway, Valerie P. Karuzis, Stefanie E. Kuchinsky

**Author notes:** Corresponding author: Ian Phillips, PhD, National Military Audiology and Speech Pathology Center Walter Reed National Military Medical Center, The Henry Jackson Foundation for the Advancement of Military Medicine, Inc., America Building, Room 5400 4954 North Palmer Road Bethesda, MD 20889.

## Abstract

Transcutaneous auricular vagus nerve stimulation (taVNS) is a neuromodulatory technique that may have numerous potential health and human performance benefits. However, optimal stimulation parameters for maximizing taVNS efficacy are unknown. Progress is impeded by disagreement on the identification of a biomarker that reliably indexes activation of neuromodulatory systems targeted by taVNS, including the locus coeruleus-norepinephrine (LC-NE) system. Pupil size varies with LC-NE activity and is one potential taVNS biomarker that has shown inconsistent sensitivity to taVNS in prior studies. The present study examined the relationship between pupil size and taVNS using stimulation parameters that have shown promising behavioral effects in prior studies but have received comparatively little attention. Participants received 30-second trains of 50 μs taVNS pulses delivered below perceptual threshold at 300 Hz to the left external acoustic meatus (EAM) while pupil size was recorded during a pupillary light reflex task. Analysis of pupil size using generalized additive mixed modelling (GAMM) revealed a nonlinear relationship between taVNS intensity and pupil diameter. Active taVNS increased pupil size during stimulation for participants who received taVNS between 2 and approximately 4.8 mA, but not for participants who received higher intensity taVNS (up to 8.1 mA). In addition, taVNS effects persisted in subsequent blocks, mitigating decreases in pupil size over the course of the task. These findings suggest 300 Hz taVNS activates the LC-NE system when applied to the EAM, but its effects may be counteracted at higher intensities.

## 1 INTRODUCTION

Transcutaneous auricular vagus nerve stimulation (taVNS) is a promising, non-invasive technique for inducing broad neuromodulatory changes throughout the brain by applying weak electrical current to parts of the outer ear that receive vagal innervation. taVNS is a lower-risk and lower-cost alternative to invasive (implanted) vagus nerve stimulation (iVNS), which has several FDA-approved applications but requires surgery and is limited to patient populations. Research on taVNS has increased markedly in recent years due to numerous potential health and human performance applications, including but not limited to treatment for epilepsy (Bauer et al., 2016; Lampros et al., 2021), depression (Hein et al., 2013; Kong et al., 2018; Rong et al., 2016), anxiety (Burger et al., 2016, 2019), posttraumatic stress disorder (Lamb et al., 2017), chronic pain (Busch et al., 2013; Farmer et al., 2020), migraine (Barbanti et al., 2015; Straube et al., 2015), cluster headache (Gaul et al., 2016), tinnitus (Hyvärinen et al., 2015; Kreuzer et al., 2012; Yakunina & Nam, 2021), and schizophrenia (Hasan et al., 2015); as well as improving upper limb motor rehabilitation (Wu et al., 2020) and infant oral feeding (Badran et al., 2020); see Verma et al. (2021) for a systematic review and discussion of study limitations. In healthy adults, taVNS has also been linked to short-term cognitive benefits in learning and memory (Giraudier et al., 2020; Jacobs et al., 2015; Llanos et al., 2020; McHaney et al., 2023; Pandža et al., 2020; Phillips et al., 2021), executive function (Ridgewell et al., 2021), as well as improved mood and reduced anxiety (Calloway et al., 2020).

Despite its promise, use of taVNS outside of the laboratory is hindered by the lack of a reliable method for titrating stimulation to achieve the desired neuromodulatory effects. Studies of iVNS parametric manipulations indicate that its efficacy depends on stimulation intensity and frequency. Moderate intensity stimulation (0.4–0.8 mA, depending on the study) has been shown to optimize the neuroplastic (Borland et al., 2016; Morrison et al., 2019, 2020) and behavioral effects (Pruitt et al., 2021) of iVNS in animal models (for a recent summary, see Hays et al., 2023). In humans, the effect of iVNS on cognition has shown a similar intensity-dependent relationship (Clark et al., 1999; Vonck et al., 2014). Importantly, in both humans and animal models, iVNS delivered outside of this narrow intensity range is ineffective (Morrison et al., 2019, 2020) and higher intensities may even have counteractive effects (Clark et al., 1999; Helmstaedter et al., 2001; Morrison et al., 2021). Efficacy of taVNS is likely to similarly vary as a function of intensity given that common mechanisms are hypothesized for both techniques (more on this below).

Parametric studies of taVNS are necessary to establish optimal stimulation parameters; however, this endeavor is complicated by the fact that it is not possible to directly measure the amount of electrical current that reaches the auricular branch of the vagus nerve (ABVN) from stimulating electrodes placed on the skin of the outer ear. Further, estimating the amount of charge reaching vagal afferents from the stimulating electrode is not possible due to individual differences in skin impedance and cutaneous distribution among regions of the outer ear that are thought to receive vagal innervation (Butt et al., 2020; Ludwig et al., 2021; Yap et al., 2020). Titrating taVNS parameters to optimize its effectiveness across individuals thus requires a biomarker that reliably indexes activation of the hypothesized target neuromodulatory systems (Burger, D’Agostini, et al., 2020). The present study contributes to this endeavor by demonstrating a nonlinear relationship between pupil dilation—one potential taVNS biomarker that has shown inconsistent prior results—and high-frequency taVNS intensity.

### 1.1 Purported mechanisms of action

The taVNS mechanism of action is thought to be like that of iVNS, which involves triggering the broad release of neurotransmitters including norepinephrine (NE; Dorr & Debonnel, 2006; Hassert et al., 2004; Hulsey et al., 2019; Manta et al., 2009; Roosevelt et al., 2006), acetylcholine (ACh; Hulsey et al., 2016; Mridha et al., 2021), and serotonin (5-HT; Dorr & Debonnel, 2006; Hulsey et al., 2019; Manta et al., 2009). This occurs via the relay of sensory information from the auricular branch of the vagus nerve (ABVN) by the nucleus of the solitary tract (NTS) to the locus coeruleus (LC) and its projections to the raphe nuclei and throughout cortex (Badran et al., 2018; Ruffoli et al., 2011; Sclocco et al., 2019); for a recent review, see Butt et al. (2020). Given the hypothesized central role of the LC-NE system in driving taVNS effects, several biomarkers that reflect LC-NE activity have been assessed for their sensitivity to taVNS, including: concentrations of salivary alpha amylase (sAA) and cortisol; pupil size (during rest, prestimulus baseline, and stimulus-evoked change); spectral power of the electroencephalogram (EEG); intensity and latency of the P3 component of the event-related potential (ERP); heart rate variability; respiratory rate; and salivary flow rate (for a review, see Burger, D’Agostini, et al., 2020). Of these measures, using pupillometry to assess taVNS efficacy is of particular interest because changes in pupil size show a relatively tight temporal coupling to fluctuations in NE as well as ACh activity (Aston-Jones & Cohen, 2005; Joshi et al., 2016; Murphy et al., 2014; Reimer et al., 2016; Samuels & Szabadi, 2008b) and can be easily measured and analyzed in near real time at the level of the individual at relatively low cost.

Direct stimulation of the LC has been shown to cause pupil dilation in animal models (Breton-Provencher & Sur, 2019; Joshi et al., 2016; Liu et al., 2017; Megemont et al., 2022; Reimer et al., 2016). Crucially, direct stimulation of vagal afferents via *iVNS* has shown similar causal effects on pupil dilation in patients (Desbeaumes Jodoin et al., 2015; but cf. Schevernels et al., 2016) and in animal models (Bianca & Komisaruk, 2007; Mridha et al., 2021), demonstrating that the impact of electrical vagus stimulation on LC activity can in principle be measured in real time via changes in pupil size. However, the impact of *taVNS* on pupillary measures has been inconsistent (Burger, D’Agostini, et al., 2020).

### 1.2 Overview of previous studies of the impact of taVNS on the pupil response

To date, 16 studies have been published examining the effects of taVNS on the size and timing of the pupil response. These studies, summarized in Table 1, have targeted three regions of the external ear known to receive vagal innervation: the concha, tragus, and external acoustic meatus (EAM; Butt et al., 2020). Thirteen of the 16 published studies targeted the concha with 200–300μs taVNS pulses delivered at 25–30 Hz (Borges et al., 2021; Burger, Van der Does, et al., 2020; D’Agostini et al., 2021, 2022, 2023; Keute et al., 2019; Lloyd et al., 2023; McHaney et al., 2023; Sharon et al., 2021; Skora et al., 2024; Urbin et al., 2021; Warren et al., 2019; Wienke et al., 2023). Of these studies, two also tested parameters that deviated from these ranges: Urbin et al. (2021) included 300 Hz taVNS and D’Agostini et al. (2023) included a 400 µs pulse-width condition. Across these studies, taVNS intensity and titration method varied. Four studies tested fixed taVNS intensity, at 0.2 and 0.5 mA (D’Agostini et al., 2023), 0.5 mA (Burger, Van der Does, et al., 2020; Warren et al., 2019), and 3.0 mA (Keute et al., 2019). Five studies tested taVNS delivered just below pain threshold, with reported intensities ranging from 0.25–5.0 mA (D’Agostini et al., 2021, 2022, 2023; Lloyd et al., 2023; Sharon et al., 2021). The remaining five studies tested taVNS intensities below, at, or above perceptual threshold (but below pain threshold), with reported values ranging from 0.08–4.4 mA (Borges et al., 2021; McHaney et al., 2022; Skora et al., 2024; Urbin et al., 2021; Wienke et al., 2023). Only six of the 13 studies targeting the concha noted an effect of taVNS on pupil size. In five studies, taVNS increased resting-state pupil size compared to a control condition (D’Agostini et al., 2023; Lloyd et al., 2023; Sharon et al., 2021; Skora et al., 2024; Urbin et al., 2021); in the sixth study, taVNS increased amplitude of the task-evoked pupillary response (TEPR) during an active task and reduced constriction during a pupillary light reflex task.

**Table 1:**
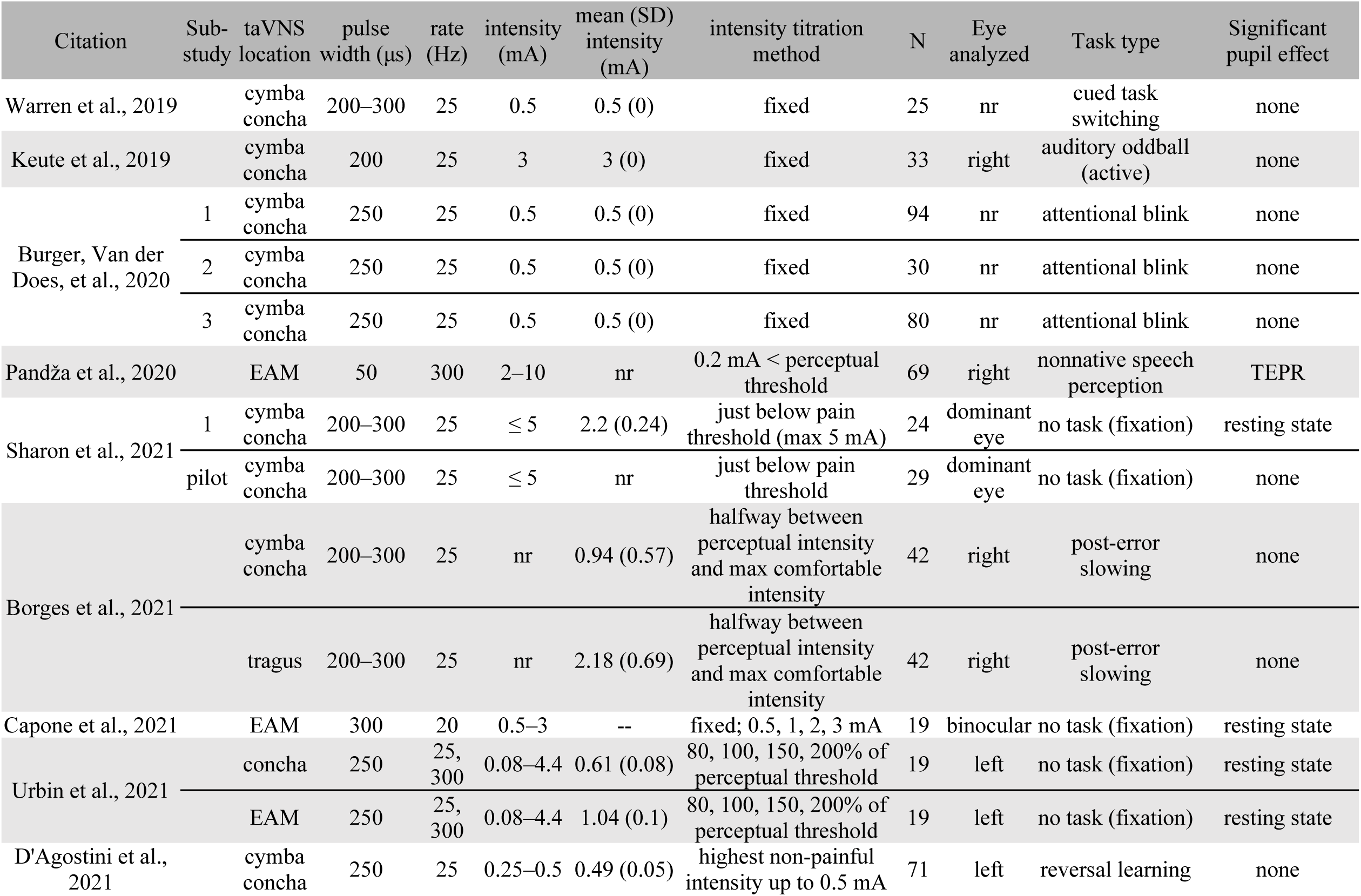
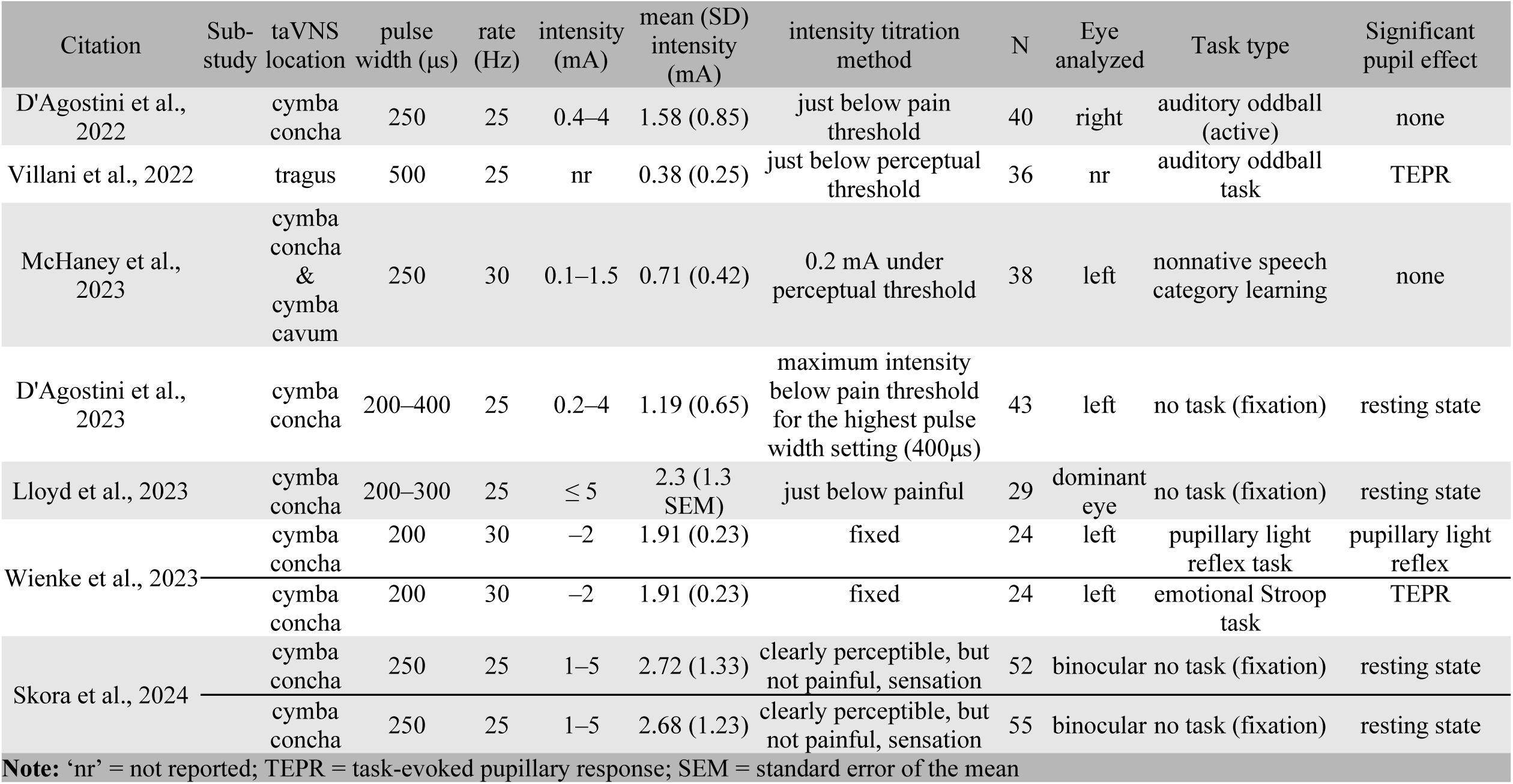
Summary of taVNS studies measuring pupillary response.

Two studies measured pupillary responses to taVNS applied to the tragus (Borges et al., 2021; Villani et al., 2022). Only Villani et al. (2022) found an effect of taVNS on TEPR amplitude, though opposite from the expected direction. In that study, 500 μs taVNS pulses were delivered at 25 Hz just below perceptual threshold for 3 seconds around stimulus onset in an auditory oddball task. The effects of taVNS were largely null, only decreasing TEPR peak amplitude for trials with low baseline pupil size determined by a median split.

Three studies have measured pupillary responses to taVNS applied to the EAM. All three studies found an effect of taVNS on pupil size. In Capone et al. (2021), 300 μs taVNS pulses delivered at 20 Hz elicited larger resting state pupil size in the left eye (but not the right eye), but only for 2.0 mA taVNS during low illuminance. No taVNS-related effects were observed under medium or high illuminance for any taVNS intensity (0, 0.5, 1.0, or 3.0 mA). Urbin et al. (2021) directly compared taVNS rate (25 versus 300 Hz), stimulation site (concha vs. EAM), and intensity (0, 80, 100, 150, and 200% of perceptual threshold). Increasing taVNS intensity generally increased resting-state pupil size and decreased onset and peak latency of the pupillary response, with the strongest effects observed for 300 Hz taVNS delivered to the EAM. taVNS delivered to the EAM at and above perceptual threshold increased pupil size over earlobe (active sham) stimulation and increased pupil diameter over concha stimulation at 150% of perceptual threshold. The area under the curve between pupil response onset and peak also showed that 300 Hz taVNS applied to the EAM elicited larger pupil responses than 25 Hz taVNS delivered to the same location and 300 Hz taVNS delivered to the concha or earlobe. Pupil response onset latency was also shorter for 300 Hz versus 25 Hz taVNS across sites, and onset latency was shorter for EAM than concha.

Pandža et al. (2020) examined behavioral and pupillary effects of delivering 50 μs taVNS pulses to the EAM at 300 Hz and 0.2 mA below perceptual threshold during a foreign language learning study. Behavioral improvements and electrophysiological changes (reported in Phillips et al., 2021) were observed for participants who received active taVNS, and these changes coincided with distinct changes in TEPR intensity across two training days. However, given the nature of the training task, the TEPR changes in Pandža et al. (2020) were interpreted as reflecting between-group differences in the allocation of effort during learning, rather than reflecting the direct effect of taVNS on the LC-NE system.

That most studies to date have found no effect of taVNS on pupil size has led to questioning whether taVNS can actually modulate LC activity (Burger & Verkuil, 2018). However, there are at least three factors that may contribute to the general lack of taVNS effects on pupil size in previous work that do not necessarily depend on taVNS failing to activate the LC. The first pertains to taVNS parameters. The taVNS parameters most often used in previous studies may not drive sufficient change in LC activity required for coupling taVNS and pupil dilation, given recent evidence that pupil dilation may not accurately track LC activity below a certain LC firing rate (Megemont et al., 2022). The second pertains to statistical analysis. Inconsistencies in the amount of charge reaching vagal afferents across participants within a study due to individual differences in skin impedance, discomfort tolerance (when suprathreshold stimulation is used), electrode placement, and cutaneous distribution of the ABVN may reduce the power to link fixed taVNS intensities to outcome measures. In studies where taVNS intensity is titrated individually based on sensory percepts, the relationship between taVNS intensity and resulting physiological changes across participants may be nonmonotonic. Such a relationship may not be captured in linear analyses. The third pertains to experimental design. Characterizing the relationship between taVNS and pupillary measures during a task where learning may occur poses potential problems. One goal of pairing taVNS with cognitive training is to reduce the difficulty or amount of cognitive effort required to sustain levels of task performance. Pupil size is known to reflect cognitive load, with less difficult or effortful tasks eliciting smaller task-evoked pupil responses (Beatty, 1982). Thus, the effect of taVNS reducing the cognitive effort required during task performance may counteract expected increases in TEPR intensity due to taVNS modulating LC activity.

### 1.3 Goals of the current study

With the goal of further evaluating the use of pupil dilation as a physiological biomarker of taVNS-related changes in LC-NE activity, the present study used a single-blind, sham-controlled, within-subjects experimental design to measure taVNS-related changes in pupil diameter during a pupillary light reflex task, similar to that used in a prior study that shows pupillary response to iVNS (Desbeaumes Jodoin et al., 2015). We selected this task in order to replicate methods that had shown an effect of VNS on pupil size as closely as possible. We addressed each of the potential issues noted above by using taVNS parameters that have shown a more robust pupillary response in prior studies (300 Hz applied to the EAM), measuring pupil changes during a passive task, and analyzing the effect of individually-titrated taVNS intensity using generalized additive mixed models (GAMMs; Wood, 2006). GAMMs allow for the characterization of nonlinear effects of taVNS intensity on the size and timing of pupillary responses while reducing Type 1 error by simultaneously modeling participant-level random effects and accounting for the high degree of autocorrelation in pupil data (van Rij et al., 2019).

Note that this study is not an exploration of taVNS parameter manipulations within participants. Rather, it aims to characterize the relationship between pupil size and a single taVNS intensity across participants. Based on the results of Desbeaumes Jodoin et al. (2015), we hypothesized a sustained increase in pupil size during trials during which participants received active taVNS compared to pupil size when they received passive sham taVNS; we did not hypothesize that active taVNS would have differential effects on components of the pupillary light reflex that reflect sympathetic versus parasympathetic influences (Samuels & Szabadi, 2008b).

## 2 METHOD

This study was approved by the Institutional Review Board (IRB) of the University of Maryland and the U.S. Department of Navy Human Research Protection Program.

### 2.1 Participants

One hundred and twelve healthy adults consented to participate in this study. Participants were 18– 31 years old (*M* = 20.46*, SD* = 2.49) and 63.5 % identified as female. The relevant task was administered as part of a larger, multisession study assessing the influence of taVNS on Mandarin lexical tone learning. Sample size was determined based on the expected effect sizes for the behavioral learning tasks administered in subsequent sessions, based in previous work by the authors (Pandža et al., 2020; Phillips et al., 2021).

Participants enrolled in this study reported having normal or corrected-to-normal vision, normal hearing, and no ear canal blockage. Individuals were excluded from participating if they were pregnant or nursing, or if they reported a history of any of the following: learning disabilities, neurological, neuropsychiatric, or psychiatric disorders; ocular disorders affecting pupillometry (e.g., cataract, nystagmus, amblyopia); recently receiving VNS/TENS outside of lab protocols; electroshock therapy for depression; taVNS contraindications including cardiac or vascular disease, diabetes, epilepsy, fainting; head or face injuries, pain, or pain disorders; implanted metallic or electronic devices, non-removable metal piercings around the face; heart attack, stroke/TIA, congestive heart failure, coronary heart disease, peripheral vascular disease, angina pectoris, irregular heartbeat or arrhythmia; concussion or loss of consciousness for more than 10 minutes; recent hospitalization; taking psychoactive medications or medications to treat taVNS contraindications within 2 months.

### 2.2 Pupillary light reflex task

The experimental task administered in this study was designed to closely follow the pupillary light reflex task in Desbeaumes Jodoin et al. (2015) used to measure changes in pupil size during iVNS in patients with refractory epilepsy or depression. The sequence of visual stimuli during each trial is shown in Figure 1. Each trial began with a 1 cm gray fixation cross shown centered over a black background on the participant computer display (Dell 24-inch LED monitor). The fixation cross was visible for 5 sec, after which the fixation cross disappeared and the black background remained unchanged for 1.5 sec. Next, a white 10 cm diameter disk appeared over the black background in the center of the display. After 0.5 sec, the disk disappeared and the black background was displayed for 1.95 sec, followed by the presentation of the gray fixation cross in the center of the screen for another 5 sec.

**Figure 1:**
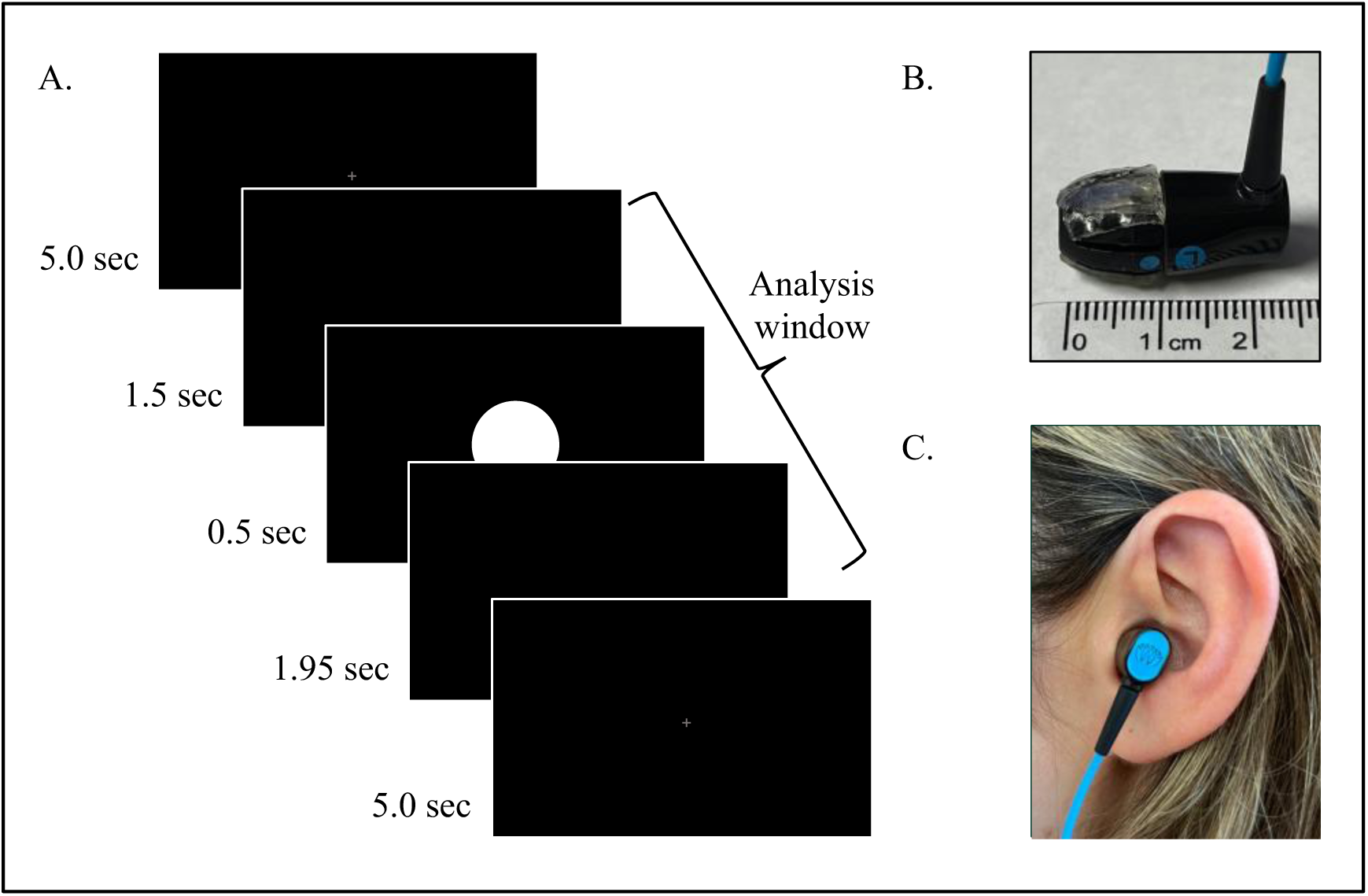
(A) Pupillary light reflex trial sequence, with timing and analysis window noted. (B) Modified earbuds used to deliver taVNS to the left EAM. Note, conductive gel was placed on the top (not shown) and bottom (shown) of the earbud, over the conductive surfaces. (C) Position of the modified earbud within the left EAM.

### 2.3 taVNS parameters and titration

A Digitimer DS8R Biphasic Constant Current Stimulator (DS8R; Digitimer North America, LLC, Fort Lauderdale, FL) generated the taVNS. Each stimulus train consisted of 50 µs biphasic square pulses with 350 microsecond interphase dwell and 100% recovery phase ratio delivered at 300 Hz. We delivered taVNS at 300 Hz, rather than the 25–30 Hz rate standardly used in taVNS studies, because of evidence suggesting it was better tolerated by participants (Tyler et al., 2019) and positive effects of these taVNS parameters were found for behavioral, pupillary, and ERP results in a previous study conducted by the authors (Pandža et al., 2020; Phillips et al., 2021). The DS8R output a single stimulus when triggered via a TTL pulse over a BNC jack, which originated from a custom-programmed Arduino UNO board controlled by the experiment computer via USB. The experiment software controlled the 300 Hz pulse delivery rate via the USB connection and also controlled the DS8R’s current intensity via a DS8R API over a secondary USB connection.

A set of modified Neuvana earbuds (previously Nervana; Neuvana, LLC, Deerfield Beach, FL), was used to deliver taVNS via a 2.5 mm electrode plug connected to the DS8R, and also delivered auditory stimuli via a 3.5 mm audio plug connected to the experiment computer. The earbuds contained two electrodes on the left earbud, positioned to stimulate the ABVN from within the EAM. The modification involved replacing a small portion of the silicone earbud tip over each electrode with Axelgaard AG735 and/or AG2550 hydrogel (Axelgaard Manufacturing Co., Ltd.) and affixing a small patch (approximately 0.5–1 cm^2^) of the same hydrogel on the surface of the earbud tip over each replaced section (see Figure 1). Hydrogel was used as the transmission medium to maintain consistent connectivity throughout the experiment.

Each participant’s taVNS perceptual threshold was determined using the titration procedure described in Pandža et al. (2020). Briefly, 2 sec tVNS pulse trains were delivered at random intervals from 1–3 sec, starting at 2 mA and increasing in 0.5 mA steps to a maximum of 10 mA. When participants pressed a button to indicate that they felt stimulation, taVNS intensity was reduced by 1.0 mA (to a minimum of 2.0 mA) and the staircase procedure was repeated in 0.1 mA steps until participants again pressed a button to indicate they felt stimulation. The intensity when the participants pressed the button the second time was taken as the perceptual threshold. During the task, taVNS was delivered 0.2 mA below perceptual threshold to maintain participant blindness to the taVNS condition (active versus sham) during each experimental block. Passive sham taVNS was used as the control condition in this study. During sham taVNS blocks, no current was delivered to either earbud.

### 2.4 Procedure

For all but one participant included in the analysis, the pupillary light reflex task was administered at the end of the first testing session of the larger study and was participants’ first encounter with taVNS in this study.^1^ The session lasted approximately 2 hours. Before the pupillary light reflex task, each participant provided informed consent and completed several questionnaires and experimental tasks to assess Mandarin lexical tone identification and categorization ability, in line with the aims of the larger study. Participants completed the pupillary light reflex task seated in front of the computer display positioned approximately 65 cm from participants’ eyes, with their heads stabilized using a chinrest. Pupil size and gaze position were recorded with a Tobii X120 eye-tracker affixed below the display, with pupil variables sampled at 40 Hz and gaze variables sampled at 120 Hz. Visual stimuli, taVNS delivery, and recording of eye data were all controlled using a custom script developed using PsychoPy (v. 3.1.5; Peirce et al., 2019).

The pupillary light reflex task consisted of 12 trials split evenly over four blocks. Each block lasted about 42 seconds with a three-minute rest period between blocks. All participants received passive sham taVNS during the first and third blocks and active taVNS continuously throughout the second and fourth blocks. At the beginning of the task, participants read task instructions on the display and then completed the taVNS threshold procedure described above followed by a nine-point eye-tracker calibration. The task instructions informed participants that they would see a series of shapes flash on the screen while their pupils are recorded and that they may receive taVNS at different points during the task. Participants were instructed to keep their chin in the chinrest throughout the task, focus on the center of the screen, limit their blinks only to when the fixation cross was on screen, and to avoid other movements.

Additional measures beyond delivering taVNS below perceptual threshold were implemented to ensure that participants were unaware of when they were receiving taVNS during the task. To mask potential sound artifacts that occasionally occurred during taVNS due to leakage of current between the taVNS and audio lines within the stimulating earbud, all participants heard a pink noise sound mask described in Phillips et al. (2021) continuously during any period when taVNS was possible, including calibration and each task block.

At the onset of the study, the testing booth was illuminated by a single lamp positioned behind the participant resulting in a luminance of 40 lux as measured at the participant’s eye (i.e., at the chinrest facing the screen). A check of data quality performed halfway through data collection indicated noisier-than-expected pupil tracking. It was determined that this was due to insufficient ambient luminance. As a result, the overhead light inside of the booth was turned on during the task for the remainder of participants, increasing luminance to 159 lux as measured at the participant’s eye.

### 2.5 Pupillometry preprocessing and analysis

Prior to analysis, pupillary data for left and right eyes were preprocessed using a custom script in R (R Core Team, 2022) based on recommendations of Kret and Sjak-Shie (2019). For each eye, pupillary data were first upsampled to the 120 Hz gaze sampling rate by linearly interpolating between valid pupil samples. This was done to improve correction of artifacts described below. Next, pupil samples were marked as invalid and removed if the difference in diameter between that sample and either neighboring sample exceeded ten times the sample-to-sample median absolute deviation across the entire trial. After removing invalid samples, contiguous segments of invalid pupil samples exceeding 75 ms were marked as blinks and samples during the 50 ms preceding and following blinks were removed due to apparent rapid changes in pupil size during these segments due to partial occlusion of the pupil by the eyelid.

Missing samples were then replaced using linear interpolation and a smoothed vector of pupil data was created using a 125 ms symmetrical moving window average. Next, pupil samples were marked as invalid and removed if the difference in diameter between that sample and the corresponding sample in the smoothed vector exceeded 0.5 times the median absolute deviation between pupil samples and their corresponding smoothed values across the entire trial. The process of linear interpolation, smoothing, and removal of outliers was then repeated a second time. The deviation multipliers used in pre-processing were determined empirically as recommended in Kret and Sjak-Shie (2019) as those that removed obvious outlier samples without resulting in excessive removal of data. Pupil samples that were missing from one eye only were replaced based on the value of the other eye and model coefficients from regressing each eye onto the other. Finally, left and right pupil diameter was averaged at each sample, remaining missing values were linearly interpolated, and the averaged pupil vector was smoothed using a 125 ms symmetrical moving window average.

The pupillary analysis window spanned the 3.95 sec comprising the white disk and the preceding and following blank screens. Pupil diameter during the blank screens before and after stimulus that elicited the pupillary light reflex was modeled in order to permit identification of taVNS effects during both the resting state and during the pupillary light reflex, which may be differentially affected by increases in LC activity resulting from taVNS (Samuels & Szabadi, 2008b). Trials in which more than one-third of samples during the analysis window were missing for both eyes prior to interpolation were excluded (32.3% of trials). As a result, data for 6 participants were excluded entirely. An additional 6 participants who did not have at least one trial in an active taVNS block and a sham taVNS block were also excluded, as were 2 participants who lacked data due to a technical error. The final dataset consisted of 822 trials across 90 participants. The remaining trial-level vectors of pupil diameter (averaged between eyes, interpolated, and smoothed) were analyzed using GAMMs.

GAMMs are designed to fit non-linear data, particularly time-series data such as the pupillary response, and have added benefits over other forms of time-series analysis (e.g., growth curve analysis) as no *a priori* assumptions about the shape of the curve are required. Instead, non-linear patterns in the dependent variable can be fit with smooth terms for continuous predictor variables, whose shape is determined by the data and constrained by a penalty for overfitting. Smooth terms were specified as tensor product interactions to allow for the simultaneous modelling of multiple independent variables, each with its own scale and its own penalty matrix. This allows for the smooth terms to capture both non-linear changes in the pupil response across time as well as how this may change as a (non-linear) function of taVNS stimulation intensity. Additionally, GAMMs allow for the inclusion of parametric terms to model mean differences between levels of factors that may or may not interact with smooth terms. Lastly, GAMMs also allow for the inclusion of a model parameter to account for autocorrelation of model residuals, which is expected to a high degree in modeling pupillary data and may inflate Type I error rates if not accounted for (see van Rij et al., 2019).

The GAMM fit to these data included a tensor product specifying for the four-way interaction between trial time, taVNS intensity, and ordered-factor terms for taVNS condition (active vs. sham), and experiment half (first vs. second half), as well as lower order terms. A term for experiment half was included to model potential changes in pupil size that persist after delivery of active taVNS during the second block. Two additional non-interacting predictors were also included: a smooth for overall trial number (1–12) to account for the expected decrease in pupil size across the course of the experiment (McLaughlin et al., 2022); and a smooth for x- and y-gaze position, to account for systematic artifactual changes in pupil size due to changes in gaze position (Brisson et al., 2013; Gagl et al., 2011).

## 3 RESULTS

Data for *N* = 90 participants were included in the analyses. Of the 112 consented participants, five were dismissed after determining during screening that they did not meet eligibility criteria for the larger study. One additional participant was dismissed because they were able to feel the taVNS at the minimum calibration intensity of 2.2 mA and two additional participants withdrew during taVNS calibration. Data for an additional 14 participants were excluded due to insufficient pupillary data (see section 2.5).

### 3.1 taVNS intensity

For the participants included in the analyses, taVNS intensity (0.2 mA below perceptual threshold) ranged from 2.0–8.1 mA (M (SD) = 3.63 (1.37)). The distribution of taVNS intensity is shown in Figure 3A. taVNS intensity was compared between participants who completed the task before and after the change in ambient lighting halfway through data collection to ensure that there was no relationship between taVNS intensity and light level, which may confound the results. Welch’s independent t-test confirms that taVNS intensity does not differ between participants who completed the task under lower ambient luminance (*M (SD)* = 3.45 (1.20) mA) and participants who competed the task under higher ambient luminance conditions (*M (SD)* = 3.87 (1.56) mA) participants, *t*(66.76) = −1.36, *p* = .176. Further, a two-sample Kolmogorov–Smirnov test also indicates that the distributions of taVNS intensity do not differ between these participant groups (*D* = 0.18, *p* = .359).

### 3.2 Pupil diameter

The GAMM was constructed to determine (1) whether pupil diameter during any part of the pupillary light reflex task differed between blocks during which active versus sham taVNS was continuously delivered, (2) whether any taVNS-block-level differences were modulated by taVNS intensity, and (3) whether any changes in pupil size during active taVNS persisted beyond the stimulating period. In the initial model, the highest-order interaction (trial time by taVNS intensity by taVNS condition by experiment half) was not significant. To facilitate interpretation, this term was removed and the GAMM was refit to the data. The summary of the resulting GAMM is shown in Table 2. Model-predicted values for pupil diameter as a function of trial time and taVNS intensity are plotted for each block in Figure 2.

**Figure 2:**
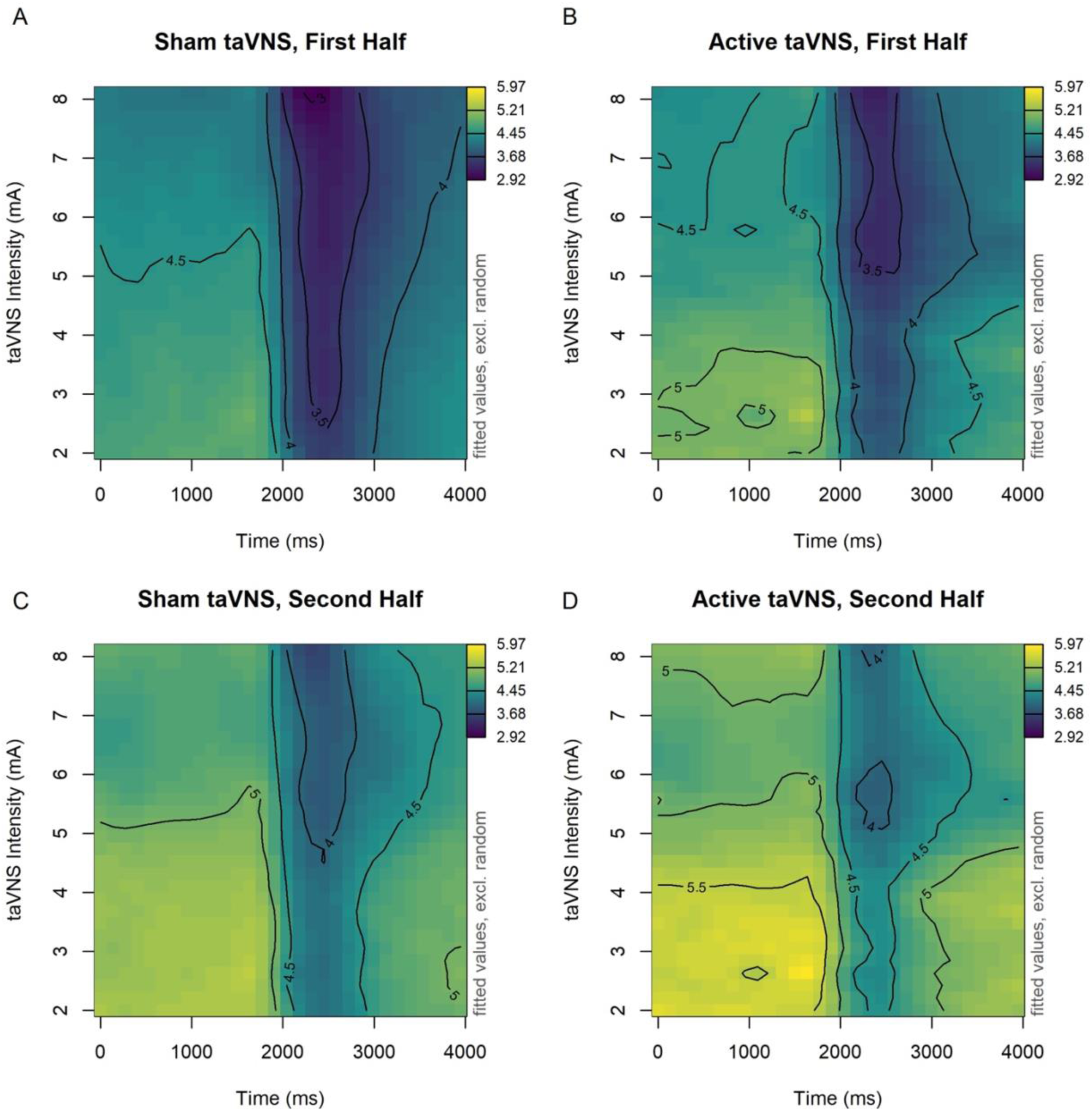
Model-predicted pupil diameter over the course of the analysis window for each task block as a function of taVNS threshold: (A) block 1, (B) block 2, (C) block 3, (D) block 4. Pupil diameter is represented in the z-axis with color (darker = smaller pupil diameter). The isobars indicate lines of equal pupil diameter in 0.5 mm steps. As a reference, the white disk was present on the display between 1500– 2000 ms.

**Table 2:** GAMM summary.

| Parametric Coefficients | Estimate | SE | t-value | p-value |
| --- | --- | --- | --- | --- |
| (Intercept) | 4.436 | 0.106 | 42.056 | < .001 |
| IsActive | 0.229 | 0.044 | 5.165 | < .001 |
| IsSecondHalf | 0.591 | 0.080 | 7.366 | < .001 |
| Smooth Terms | EDF | Ref. DF | F-value | p-value |
| s(trial number) | 7.356 | 7.872 | 84.867 | < .001 |
| s(x-position, y-position) | 2.004 | 2.008 | 1.632 | .195 |
| te(time, taVNS intensity) | 188.997 | 214.451 | 51.313 | < .001 |
| te(time, taVNS intensity):IsActive | 226.367 | 285.039 | 2.231 | < .001 |
| te(time, taVNS intensity):IsSecondHalf | 152.860 | 209.123 | 1.544 | < .001 |
| Random Smooth Terms | EDF | Ref. DF | F-value | p-value |
| s(time, subject) | 379.837 | 443.000 | 31.436 | < .001 |
| s(time, subject):IsActive | 232.775 | 438.000 | 2.725 | < .001 |
| s(time, subject):IsSecondHalf | 272.791 | 425.000 | 4.577 | < .001 |

*Note:* Model formula: pupil\_size ~ IsActive + IsSecondHalf + s(overalltrial) + s(Xpos, Ypos) + te(time, taVNS intensity, k = 20) + te(time, taVNS intensity, by = IsActive, k = 20) + te(time, taVNS intensity, by = IsSecondHalf, k = 20) + s(time, subject, bs = "fs", m = 1, k = 5) + s(time, subject, by = IsActive, bs = "fs", m = 1, k = 5) + s(time, subject, by = IsSecondHalf, bs = "fs", m = 1, k = 5). Adjusted $R^2 = 0.931$ , Deviance Explained = 81.2%, fREML = 9413.8, n = 122,138. EDF = effective degrees of freedom; Ref. DF = reference degrees of freedom.

For each significant smooth, model-predicted values are plotted to allow for interpretation, as is standard practice in GAMMs analysis (Wieling, 2018). The significant smooth *s(trial number)* indicates that pupil diameter varied as a function of trial number (*p* < .001). Model-predicted values for this effect are plotted in Figure 3B. This plot indicates a fairly linear decrease in pupil diameter across the 12 trials in the task. When accounting for this overall decrease, however, the significant parametric term *IsSecondHalf* indicates that the decrease in pupil size was significantly weaker than expected during the second half of the task compared to the first half (*Est.* = 0.591, *p* < .001). Also, the difference between task halves was not consistent across taVNS intensities, indicated by the significant tensor product smooth *te(time, taVNS threshold): IsSecondHalf* (*p* < .001). Visual inspection of this tensor product, plotted in Figure 4, reveals that pupil diameter during the second half of the task was larger than expected across the analysis window for all participants except those who received taVNS between approximately 6.0–7.0 mA. For these participants, the effect was significant only after the first 500 ms of the analysis window.

**Figure 3:**
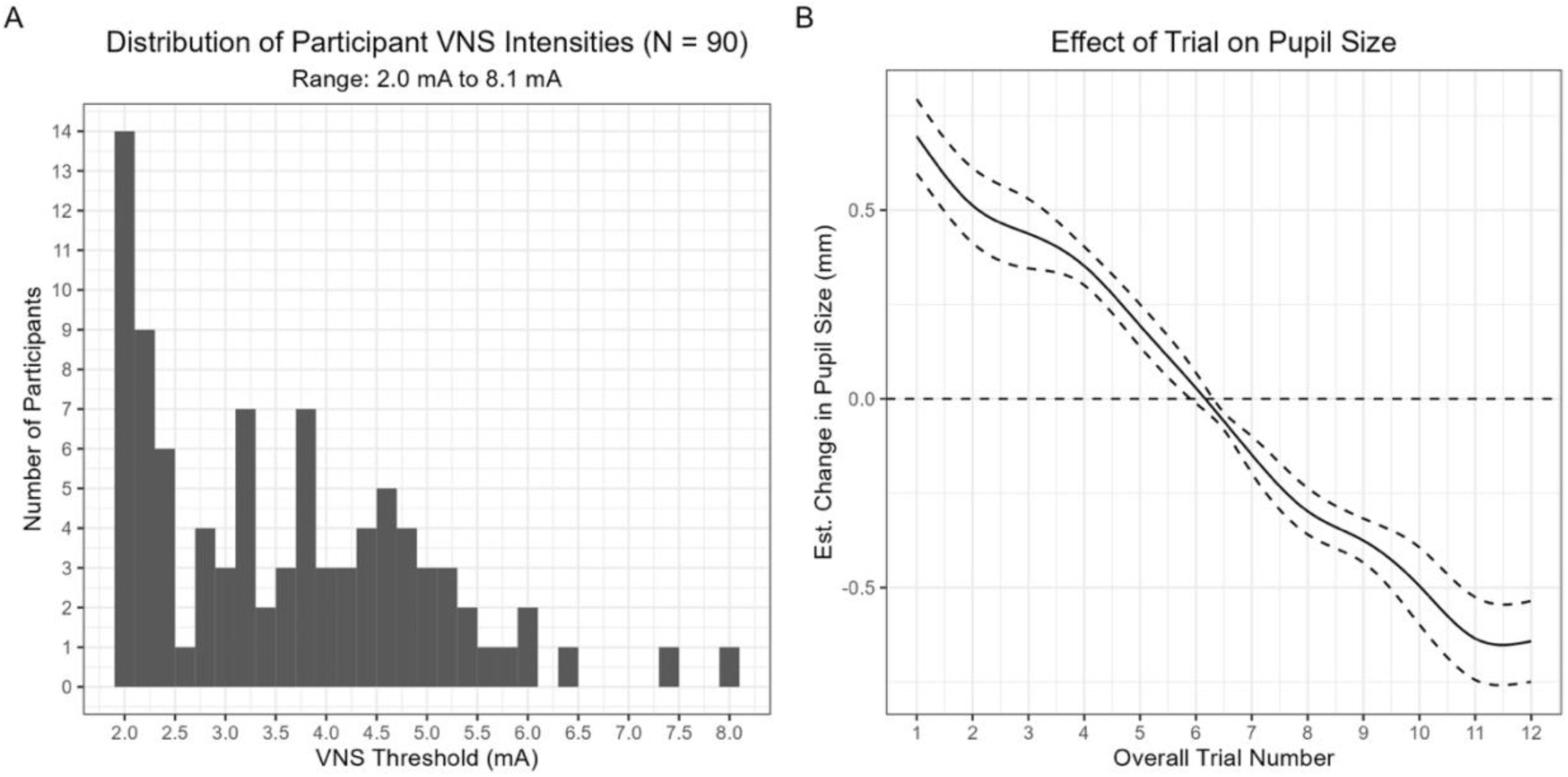
(A) Histogram of taVNS intensity levels across participants. (B) Model-predicted change in pupil diameter from grand mean (represented by the dotted line a y = 0.0) as a function of trial number.

**Figure 4:**
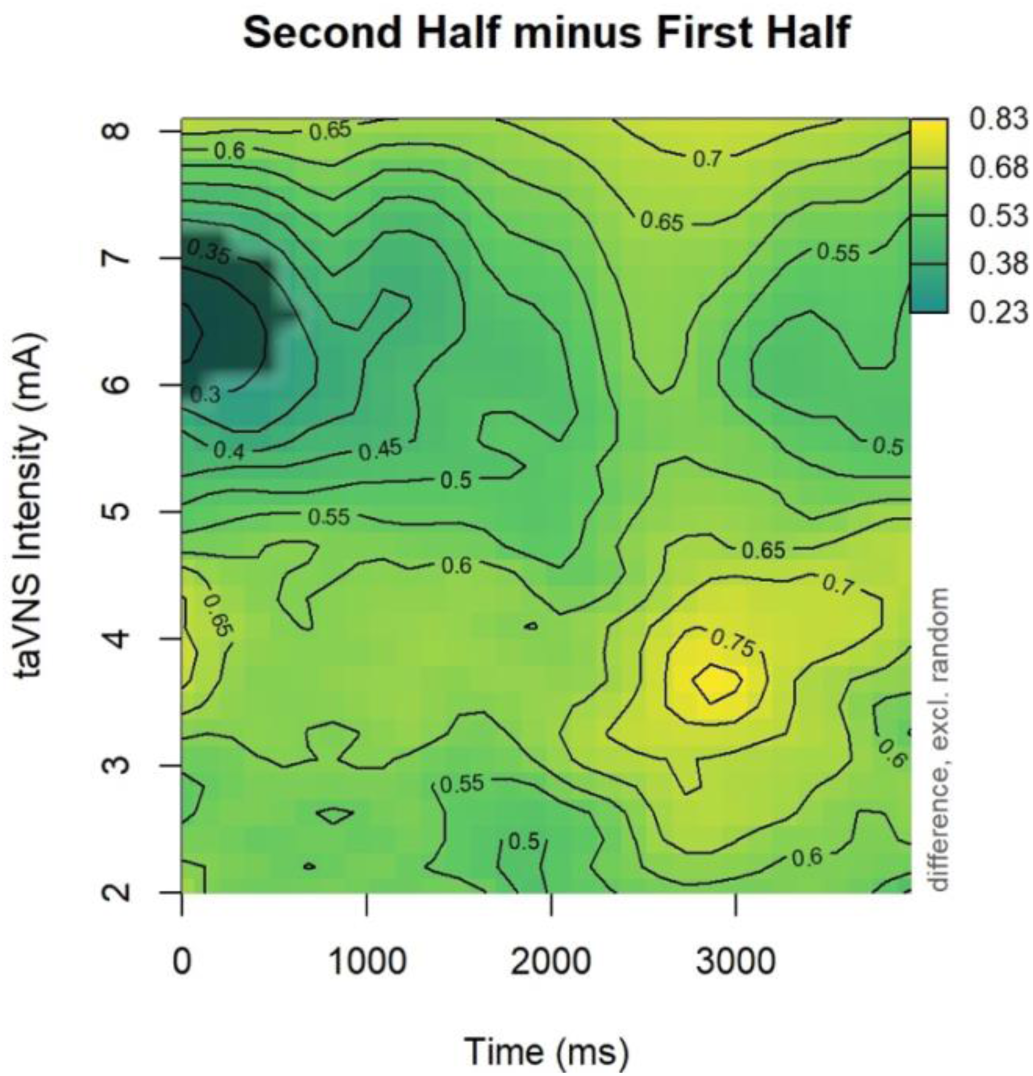
Model-predicted differences in pupil diameter between the first and second half of the task as a function of taVNS intensity. The difference in pupil diameter between task halves is indicated along the z-axis with color. Highlighted regions indicate where differences are significant.

The parametric term *IsActive* in Table 2 indicates that pupil diameter across the analysis window was significantly larger for active versus sham taVNS blocks (*Est. =* 0.229, *p* < .001). The significant tensor product smooth *te(time, taVNS threshold):IsActive* further indicates that this effect varied as a function of taVNS intensity (*p <* .001). To interpret this effect, this tensor product is plotted in Figure 5 and includes fitted smooths at the 25^th^, 50^th^, and 75^th^ percentile taVNS intensity values. This plot reveals that the increase in pupil diameter for active versus sham taVNS was significant for participants with taVNS intensities between 2 and ∼4.8 mA, but not for participants with higher taVNS intensities, specifically between ∼4.9 and 8.1 mA.

**Figure 5:**
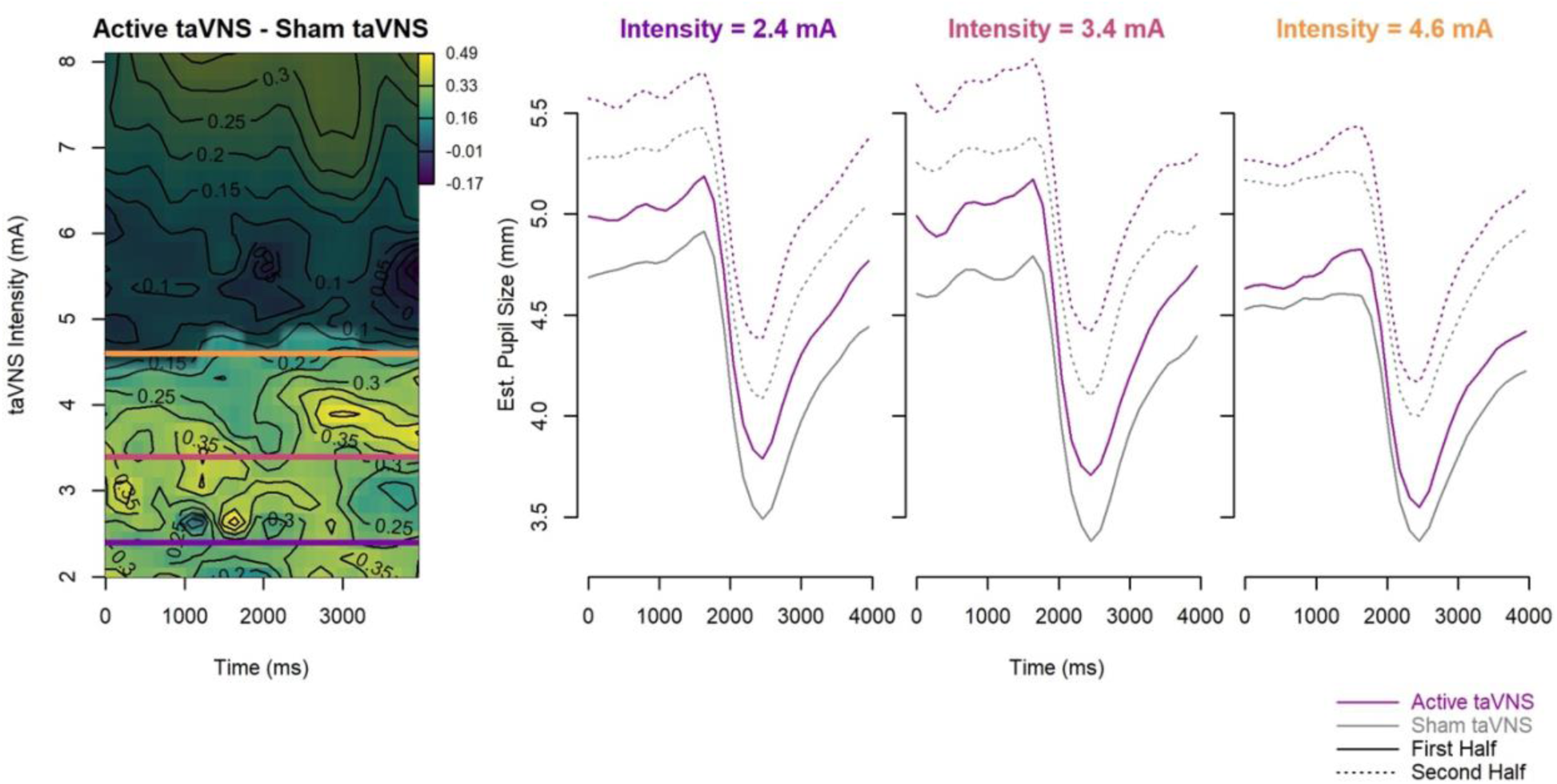
Model-predicted differences in pupil diameter between active and sham taVNS as a function of taVNS intensity. The heatmap represents the difference between the active and sham taVNS conditions as a function of time (x-axis) and taVNS intensity (y-axis). The estimated difference in pupil diameter between the two conditions is indicated along the z-axis with color. Highlighted regions indicate a significant difference between the two conditions. Horizontal lines represent the 25^th^, 50^th^, and 75^th^ percentile intensity values, for which fitted smooths are plotted in the three panels to the right. Note: Separate lines are plotted for taVNS condition and experiment half, even though this interaction was not specified in the best-fitting model because it is not possible to average over non-interacting factors when plotting GAMM-predicted values. As a reference, the white disk was present on the display between 1500–2000 ms.

## 4 DISCUSSION

The present study analyzed the effects of 30-second trains of taVNS on pupil diameter during a pupillary light reflex task in order to further evaluate the use of pupil dilation as a biomarker of taVNS-related changes in LC activity. The main result of this study indicates that delivering sub-perceptual threshold taVNS (50 µA pulses at 300 Hz) to the left EAM temporarily increased pupil size for intensities between 2.0 and ∼4.8 mA, but not higher intensities (up to 8.1 mA). A secondary finding was that taVNS effects on pupil dilation persisted across experimental blocks and this effect was relatively consistent across taVNS intensities.

### 4.1 Transient effects of taVNS on pupil size

The model-estimated effect of taVNS on pupil diameter during stimulation in the present study was an increase of 0.229 mm. This effect is very close to the average increase in resting pupil diameter of 0.23 mm elicited by iVNS using a nearly identical task in Desbeaumes Jodoin et al. (2015). Thus, our findings for taVNS appear to replicate and extend the effect of iVNS on pupil size found in Desbeaumes Jodoin et al. (2015) and the effects of taVNS on pupil size in Wienke et al. (2023), albeit with different VNS stimulation parameters. Our finding of increased pupil size with active taVNS across the entirety of the analysis window generally align with those reported in Wienke et al. (2023), in which 200 μs taVNS pulse delivered at 30 Hz at or just below 2.0 mA to the cymba concha attenuated pupil constriction during most of the analysis window (100–1200 ms, 1900–7000 ms) following onset of stimulus eliciting the light reflex, although Wienke et al. (2023) crucially differed from the present study in that taVNS was delivered for only 500 ms starting at the onset of the light reflex stimulus. In Desbeaumes Jodoin et al. (2015), 250–500 µs iVNS pulses were delivered at 20–30 Hz between 0.25–2.25 mA. However, it should be noted that the significant effect of iVNS found in Desbeaumes Jodoin et al. (2015) was based on resting pupil diameter during the 1 second prestimulus baseline period preceding presentation of the white disk, although the plot of the effect in the paper suggests it is maintained over the entire 3.95 sec trial (Desbeaumes Jodoin et al., 2015, fig. 1). We did not isolate the prestimulus period in the analysis of our pupillary data. Instead, we modeled the entire pupil time series in order to characterize taVNS effects over the course of the pupillary light reflex and the preceding resting-state period.

The model-estimated interaction between taVNS intensity and condition (active vs. sham) also suggests a non-linear effect of sub-perceptual taVNS intensity on resting-state pupil dilation. Pupil diameter was significantly larger during active versus sham taVNS for participants who received taVNS between 2 and ∼4.8 mA, but the difference in diameter was not significant for participants who received taVNS at higher intensity levels. This is of particular interest because of apparent inconsistencies in the relationship between VNS intensity and changes in pupil size noted in prior studies. Importantly, the expected relationship between VNS intensity, LC activity, and pupil size may vary depending on the nature of the experimental task and the pupil analysis window. For example, pupil dilation evoked by a task stimulus, such as appearance of a target during a target detection task, is thought to reflect LC phasic activity (Gilzenrat et al., 2010; Murphy et al., 2014). During active tasks (i.e., those that elicit behavioral responses to stimuli), the magnitude of the task-evoked pupil response (TEPR) would be expected to be maximal at moderate levels of LC activity, given the nonmonotonic (inverted-U shaped) relationship between LC phasic and tonic activity (Aston-Jones & Cohen, 2005). Phasic peak pupil dilation following VNS onset may likewise be expected to vary as a function of prestimulus baseline pupil size (Johns et al., in press; Relaño-Iborra et al., 2022), which reflects tonic LC activity (Mridha et al., 2021). However, a *monotonic* relationship between pupil dilation and LC activity might be expected for passive tasks, given evidence from mice that shows pupil dilation to increase monotonically with LC spiking rate (Megemont et al., 2022) as well as iVNS intensity, pulse width, and stimulation rate (Mridha et al., 2021).

In humans, the relationship between VNS intensity and pupil dilation has been inconsistent. Desbeaumes Jodoin et al. (2015) did not find a correlation between iVNS intensity and any pupil size metric derived from the light-reflex in humans, which the authors suggested may be due to low sample size and restricted range of iVNS intensities. In contrast, Vespa et al. (2022) found a nonmonotonic, inverted-U shaped relationship between absolute iVNS intensity and magnitude of the late pupil dilation response (PDR; 2.5–5 s following iVNS onset) in epileptic patients. Notably, the relationship between absolute iVNS intensity and the early PDR (0–2.5 s following iVNS onset) was sigmoidal. Using taVNS, Urbin et al. (2021) found the magnitude and incidence of pupillary dilation in humans increased monotonically with increasing taVNS stimulation intensity at and above (up to two times) perceptual threshold (0.08–4.4 mA; Urbin et al., 2021); and D’Agostini et al. (2023) found a linear relationship between charge per pulse (intensity x pulse width) and pupillary dilation and its temporal derivative. However, the effects in D’Agostini et al. (2023) disappeared or were drastically reduced when the highest intensity (just below pain threshold) was excluded from the analysis. Wienke et al. (2023) found a relationship between increasing taVNS intensity across a fairly narrow range up to 2 mA and reduced pupil constriction during 300–6500 ms following onset of the pupillary light reflex stimulus. More similar to the present study, Capone et al. (2021) found that moderate intensity (2 mA) taVNS, but not higher or lower intensities, elicited a pupil response.

There are several possible factors contributing to the lack of pupil dilation for active taVNS delivered above ∼4.8 mA relative to sham taVNS in the present study. One possible explanation involves activation of other pathways that may increase pupil constriction or inhibit pupil dilation. For example, activation of LC-GABA neurons may inhibit activity of LC-NE neurons and, consequently, pupil dilation (Breton-Provencher & Sur, 2019). Off target activation of these neurons has been cited as a potential explanation for the lack of strong correlation between LC activity and pupil diameter in previous VNS studies (Burger, D’Agostini, et al., 2020; Capone et al., 2021).

A similar explanation involving off-target nerve activation may also help explain the monotonic relationship between VNS intensity and LC activity found in prior studies. One possibility is that increases in taVNS intensity above perceptual threshold in prior studies may potentially activate other nerves via spreading which could result in a coupling of intensity and pupil size that does not depend on activation of the ABVN. For example, Mridha et al. (2021) found that the highest VNS intensity in a study delivering iVNS to mice elicited pupil dilation even after the vagus was resected. Although the anatomy of mice and humans is very different, this mechanism may be relevant to taVNS. Perhaps delivering taVNS below perceptual threshold, as in the present study, is less likely to cause spreading, which could result in a functionally distinct pupil response to increasing taVNS intensity that reflects primarily, if not solely, ABVN activation.

### 4.2 Persistent effects of taVNS on pupil size

The second finding is that the increase in pupil size due to active taVNS in block 2 persisted into the second half of the task (blocks 3 and 4), mitigating an overall decrease in pupil size across trials. Pupil size often decreases over the course of an experimental task (McLaughlin et al., 2022). Our finding that pupil size generally decreased over the course of our light-reflex task is in line with this. Countering this general decrease in pupil across the 12 trials, we found that pupil size during the second half of the task was larger than expected for all but a few participants, and for those participants only during a short portion at the beginning of the trial. This effect suggests a carryover effect of taVNS across blocks.

The longevity of taVNS effects is not consistent across studies, but there is some evidence that short trains of VNS can increase NE levels for up to 80 minutes in rats (Hassert et al., 2004). This provides some support for the interpretation of carryover effects given the relatively short duration of this task. It is not clear why this effect was not significant during the first 500 ms of the analysis window for participants who received taVNS between ∼6.0–7.0 mA. However, it is important to note that this effect was significant across the analysis window for the vast majority of participants across the range of taVNS intensities tested, including those who received taVNS above ∼4.8 mA. This suggests that *all* taVNS intensities that were tested in this study led to some increase in pupil size that persisted *after* block 2 taVNS ended, although only intensities less than ∼4.8 mA significantly increased pupil size *during* taVNS. It is not obvious why taVNS intensities above ∼4.8 mA would have a delayed, but not immediate, effect on pupil size. Further study is needed to replicate and explain this finding.

### 4.3 Contributions of taVNS parameters, task, and analysis approach in finding taVNS effects

To the extent that taVNS engages the same mechanisms as iVNS, its effectiveness may similarly depend on stimulation intensity. Yet, there have been few systematic evaluations of taVNS intensity effects in humans (for exceptions see Capone et al., 2021; D’Agostini et al., 2022; Urbin et al., 2021). Studies investigating taVNS have typically tested one intensity level, which is either fixed across participants or individually titrated based on sensory thresholds and these studies have found largely null effects on biomarkers known to reflect activity in the neuromodulatory systems taVNS is thought to engage. The use of GAMMs to fit potentially nonlinear effects of taVNS intensity allowed us to find a specific range of taVNS intensities that elicited an immediate increase in pupil size without the need to limit analyses to summary measures or *a priori* determined windows of interest. Furthermore, GAMMs readily allows for modeling the potential influence of autocorrelation and gaze position on pupillary effects (van Rij et al., 2019). It is possible that fitting the same data with a linear model may have obscured the same effect. In addition, the ability to examine differences in height (via the parametric terms) and differences in trajectories (via the smooth terms) aids in determining whether shifts in the pupillary response are holistic (e.g., irrespective of time) or specific to certain time windows during the window of analysis.

The present analysis approach provides a template that can be used in future studies that seek to identify optimal taVNS parameters. The sections that follow highlight potential differences between the current study and previous studies which may explain variation in the sensitivity of the pupil response to taVNS. Future studies should aim to explore these parameter spaces by testing the impact of these dimensions on the pupil response, ideally within the same participants.

#### 4.3.1 Active versus passive tasks

The use of a passive task, only requiring the participant to fixate on a visual object during taVNS, may have also contributed to observing taVNS-related pupil dilation in this study. Seven out of nine prior studies that found taVNS effects on pupil size used a passive task that only required visual fixation (Capone et al., 2021; D’Agostini et al., 2023; Lloyd et al., 2023; Sharon et al., 2021; Skora et al., 2024; Urbin et al., 2021; Wienke et al., 2023). In comparison, the seven studies that did not find effects of taVNS on pupil size all used active behavioral tasks, which required participants to respond to stimuli and measured aspects of performance such as cued task switching (Warren et al., 2019), auditory discrimination (D’Agostini et al., 2022; Keute et al., 2019), attentional blink (Burger, Van der Does, et al., 2020), post-error slowing (Borges et al., 2021), reversal learning (D’Agostini et al., 2021), and nonnative speech category learning (McHaney et al., 2023).

Three studies that found taVNS effects on pupil size involved active behavioral tasks, including an emotional Stroop task (Wienke et al., 2023), an auditory oddball task (Villani et al., 2022), and several auditory tasks involving discrimination and recognition of foreign language speech sounds and lexical learning (Pandža et al., 2020). In two of these studies, active taVNS was linked to a reduced pupillary response (Pandža et al., 2020; Villani et al., 2019). This finding is not unexpected given that one potential use of taVNS is to facilitate learning and improve cognitive performance by increasing neuroplasticity and cognitive efficiency. The facilitative effects of taVNS could reduce the cognitive effort during task performance by increasing processing efficiency (Pihlaja et al., 2020; Ridgewell et al., 2021), which would be expected to reduce TEPR amplitude (Beatty, 1982). The lack of increased pupil size due to taVNS in experiments that involve active behavioral tasks may relate to reduction in cognitive effort during active versus sham taVNS, resulting in comparatively smaller TEPRs for active vs. sham taVNS that might cancel out effects of taVNS on LC activation that go in the opposite direction, i.e., larger pupil size for active vs. sham taVNS. Teasing these effects apart would benefit the identification of a taVNS biomarker.

#### 4.3.2 Variation in taVNS intensity

Additionally, there are several differences between the taVNS parameters used in this and in prior studies that may contribute to apparent differences in the relationship between taVNS intensity and pupil dilation. Nine of the prior 16 taVNS studies that examined effects on pupil size reported testing taVNS intensities that overlapped in part or in whole with the taVNS intensity range in the present study (Capone et al., 2021; D’Agostini et al., 2022, 2023; Keute et al., 2019; Lloyd et al., 2023; Pandža et al., 2020; Sharon et al., 2021; Skora et al., 2024; Urbin et al., 2021). Another four studies tested taVNS ranges below the minimum intensity in the present study (Burger, Van der Does, et al., 2020; D’Agostini et al., 2021; McHaney et al., 2023; Warren et al., 2019). The remaining three studies did not report taVNS ranges, but the reported taVNS intensity means and standard deviations indicate taVNS intensity was below 2.0 mA for at least some participants (Borges et al., 2021; Villani et al., 2022; Wienke et al., 2023). Ten of the twelve studies that tested overlapping taVNS intensities, but none of the four studies that tested only lower intensities, found pupil effects.

At first glance, these patterns of results might suggest that a minimum taVNS intensity level may be required to elicit a pupil response. It is not possible to evaluate this hypothesis with the present data because taVNS thresholds below 2.0 mA were not tested due to the DS8R being unable to reliably deliver lower intensities.^2^ Without knowing the effects of taVNS on pupil size below 2.0 mA, it is not possible to determine whether the efficacy of the taVNS parameters tested here remains the same or changes at lower stimulation intensities. Results of prior studies also suggest that increasing taVNS intensity alone is not necessary or sufficient to elicit a pupil response. At least some participants in Urbin et al. (2021) received active taVNS below 2.0 mA, yet significant increases in pupil size over sham were observed at both taVNS sites (concha *M(SD)* = 0.61(0.08) mA; EAM *M(SD)* = 1.04(0.1) mA; range = 0.08–4.4 mA across sites). In contrast, Keute et al. (2019) delivered 3.0 mA taVNS to all participants and found no effect on pupil size. Thus, differences in other taVNS parameters likely contribute to taVNS eliciting pupil changes.

#### 4.3.3 Variation in location and configuration of the stimulating electrode

Location and configuration of the stimulating electrode may also be important. All three experiments in past studies that applied taVNS to the EAM found pupil effects (Capone et al., 2021; Pandža et al., 2020; Urbin et al., 2021), while one of the two experiments that targeted tragus found pupil effects (Villani et al., 2019), and only eight of the 18 experiments in prior studies that targeted the concha found an effect on pupil size (D’Agostini et al., 2023; Lloyd et al., 2023; Sharon et al., 2021; Skora et al., 2024; Urbin et al., 2021; Wienke et al., 2023). This suggests that stimulating the EAM is perhaps more consistently effective in increasing neuromodulatory activity. The only study to compare taVNS delivered to the EAM and the concha within participants found that perceptual thresholds were significantly greater at the EAM than the concha and that EAM stimulation generally elicited more robust pupil effects than concha stimulation (Urbin et al., 2021). Importantly, the authors noted that larger pupil dilation found for EAM versus concha stimulation were not due to differences in absolute current applied between the sites.

It is generally accepted that the cymba concha is exclusively innervated by the ABVN, while the EAM, inner tragus, and cavum concha receive afferent innervation from the vagus as well as the greater auricular nerve and auriculotemporal nerve (see Butt et al., 2020 for a comprehensive review of the available evidence). Vagal innervation of the posterior aspect of the EAM noted in cadaver studies suggest the suitability of this region for applying taVNS. However, studies using functional magnetic resonance imaging (fMRI) largely suggest that applying taVNS to the cymba concha or inner tragus (Badran et al., 2018; Frangos et al., 2015; Yakunina et al., 2017), but not the EAM, activates brain centers that are hypothesized in the taVNS mechanism of action (Yakunina et al., 2017).

Given these imaging results, it is not clear why stimulating the EAM would more consistently elicit a pupillary response across studies. One possibility is that weak activation of NTS and LC in Yakunina et al. (2017) during EAM stimulation was because taVNS was applied to the inferoposterior wall of the EAM. Vagal innervation of the EAM may be localized to the posterosuperior wall (Kiyokawa et al., 2014), which is perhaps consistent with differential effects of applying taVNS to the posterior versus anterior EAM on activation of brain stem structures (Kraus et al., 2013). Pupil effects in the present and prior studies targeting the EAM may be due in part to stimulating electrodes contacting relatively large areas of the superior and posterior walls of the EAM (Pandža et al., 2020; Urbin et al., 2021) or overlapping with the inner side of the tragus (Capone et al., 2021). In addition to potentially engaging more vagal afferents with when using larger surface area electrodes (Pandža et al., 2020; Urbin et al., 2021), targeting superior and posterior aspects of the EAM may be more effective due to the larger number of A and B myelinated afferent fibers in these aspects of the EAM, which could potentially facilitate the central effects of taVNS applied to this location (Bermejo et al., 2017) as A and B nerve fiber activation is thought to drive therapeutic effects of VNS (Ruffoli et al., 2011). Although these afferents in the EAM likely do not belong exclusively to the ABVN, all may potentially drive increased activity of the NTS when stimulated with taVNS (Bermejo et al., 2017).

#### 4.3.4 Variation in pulse width and stimulation frequency

Another key difference is that a much shorter pulse width and higher stimulation frequency (50 μs, 300 Hz) were used in the present study compared to past taVNS studies, which almost universally tested 200–300μs taVNS pulses delivered at 25–30 Hz. The focus on 20–30 Hz stimulation comes from iVNS work, which showed 50 Hz to be damaging to the vagus (Groves & Brown, 2005). It is unclear whether the same frequency constraints apply to taVNS given the many differences from iVNS.

One study, Urbin et al. (2021) directly compared 25 to 300 Hz stimulation and found that 300 Hz taVNS elicited a shorter latency pupil response onset and peak at both the concha and EAM but the only frequency effect on pupil response intensity was that 300 Hz elicited a greater area under the curve than 25 Hz and only when applying taVNS to the EAM. Evidence from rats and mice indicates that increasing the total charge per VNS pulse, by increasing the pulse width or stimulation intensity, increases the firing rate of LC neurons (Hulsey et al., 2017) and pupil dilation (Mridha et al., 2021), but the effect plateaus above a certain total charge (Hulsey et al., 2017). When the number of VNS pulses is held constant, increasing the frequency of pulse delivery does not alter the total amount of LC activity, but it does increase the amount of LC activity per unit time (Hulsey et al., 2017).

In principle, delivering 50 µs taVNS pulses at 300 Hz in the present study could drive higher levels of LC activity per unit time than past studies, which almost universally used 200–300 µs pulses delivered at 25–30 Hz (see Table 1). Although the pulse width in the current study is one-fifth to one-sixth the length used in prior studies, the pulse rate is ten times higher. Additional support for this idea comes from findings in rats that iVNS delivered at 300 Hz and above increased the synchrony of LC neuron firing compared to iVNS at 30 Hz and below (Farrand et al., 2023). In line with this, 100 Hz taVNS delivered to the cymba concha elicited stronger activation of the NTS and LC in humans compared to 2, 10, and 25 Hz stimulation (Sclocco et al., 2020). One additional benefit of using a shorter pulse is that it may be possible to attain higher taVNS intensities while maintaining stimulation below perceptual threshold (Badran et al., 2019; Tyler et al., 2019, tbl. 1). Because pupil dilation increases with LC firing rate (Joshi et al., 2016; Megemont et al., 2022; Reimer et al., 2016), perhaps some combination of higher taVNS intensity and frequency contributed to pupil dilation in the present study.

#### 4.3.5 Variation in the nature of the sham comparison

All but one prior study of taVNS effects on pupil size included an active sham taVNS comparison, whereas passive sham taVNS was used in the present study. Although the earlobe lacks vagal innervation, stimulating the earlobe, especially when above perceptual threshold, may lead to pupil dilation, thus obscuring the effects of active stimulation applied to target locations with vagus innervation (Butt et al., 2020). Indeed, active sham taVNS applied to the earlobe has been found to elicit pupil responses in some previous studies (Urbin et al., 2021).

Relatedly, although we were careful to deliver taVNS below each participant’s perceptual threshold in the present study, it is possible that the observed increase in pupil size during active compared to sham taVNS blocks could be driven in part by sensation of the taVNS (Beatty & Lucero-Wagoner, 2000). For example, noxious stimuli can elicit pupil dilation (e.g., electrical fingertip stimulation; Chapman et al., 1999). We cannot address this possibility directly because we did not obtain information from participants about sensations they might have perceived during the pupillary light reflex task or whether they could identify the task blocks during which they received active taVNS. However, analyses of off-target taVNS effects and blinding efficacy measures that were obtained during subsequent testing sessions when taVNS was paired with training are encouraging. In the sample of participants who completed all four training sessions, which includes all but 16 participants who are represented in the present study, those who received active taVNS did not rate feeling more pain or irritation in the ear receiving taVNS or other sensations compared to participants who received passive sham taVNS during training. Further, among participants who received active taVNS during training sessions, the vast majority (68.4%–88.2% across groups) were unable to correctly identify when they received active taVNS during the training tasks.

#### 4.3.6 Analysis of the left versus right eye

One final aspect of the present analysis approach may have contributed to finding taVNS effects on pupil size. The present analysis was based on pupil data averaged from the left and right eyes. A recent study found that taVNS elicited pupil dilation in the left but not right eye, ipsilateral to the stimulation ear (Capone et al., 2021). Findings in rats also indicate that LC stimulation elicits significantly larger pupil dilation in the eye ipsilateral to stimulation and LC stimulation influences the contralateral pupil via parasympathetic pathways only, while ipsilateral pupil dilation is mediated by sympathetic and parasympathetic pathways (Bianca & Komisaruk, 2007; Liu et al., 2017).

Of the nine prior studies that found a taVNS pupil effect, three analyzed pupil data from the left eye (D’Agostini et al., 2023; Urbin et al., 2021; Wienke et al., 2023), one analyzed pupil data from the right eye (Pandža et al., 2020), two analyzed data averaged for left and right eyes (Capone et al., 2021; Skora et al., 2024), and three did not report whether analysis was based on the left or right eye (Lloyd et al., 2023; Sharon et al., 2024; Villani et al., 2022).^3^ The study that the present work was based on also found left-eye pupil dilation following iVNS (Desbeaumes Jodoin et al., 2015).

Of the seven studies that did not find a pupil effect, two analyzed left-eye data (D’Agostini et al., 2021; McHaney et al., 2023), three analyzed right-eye data (Borges et al., 2021; D’Agostini et al., 2022; Keute et al., 2019), and two did not report the analysis eye (Burger, Van der Does, et al., 2020; Warren et al., 2019). Thus, while differences in the presence of pupil effects between taVNS studies examining the left versus right pupil are somewhat inconsistent, the lateralization of pupil effects noted above suggest that analysis of left-eye pupil data may be more likely to yield robust findings in future taVNS studies given that taVNS is almost always administered to the left ear to avoid potential adverse cardiac effects (Farmer et al., 2021). However, whenever possible, recording data from both eyes offers the advantage of filling in samples that are missing from one eye. Furthermore, without a strong a priori hypothesis regarding lateralization effects, analyzing the average of the two eyes (as done in the current study) may represent a more conservative approach to examining the impact of taVNS on the pupil response. Interestingly, related to the earlier discussion of stimulation rate, Liu et al. (2017) showed that the relative contribution of the ipsilateral sympathetic pathway to pupil dilation was inversely related to stimulation rate, while there was no frequency effect for the relative contribution of the ipsilateral parasympathetic pathway.

### 4.4 Limitations and future directions

Pupil dilation reflects activity of several neural circuits including the principal neuromodulatory systems VNS is thought to activate (Joshi & Gold, 2020; Strauch et al., 2022). Given this, it is not possible to attribute taVNS-related pupil dilation in the present study specifically to LC activation (Larsen & Waters, 2018). Additionally, pupil dilation is influenced indirectly by both inhibitory and excitatory inputs from the LC. Increased LC activity can inhibit parasympathetic activity of the Edinger-Westphal nucleus (EWN), thus inhibiting pupil constriction (Samuels & Szabadi, 2008a). LC activation attenuates the pupillary light reflex via its inhibitory effects on the EWN (Samuels & Szabadi, 2008b). Increased LC activity can also increase sympathetic excitatory input to the superior cervical ganglion (SCG), thus driving pupil dilation (Mridha et al., 2021).

Measuring pupil dilation preceding and during the pupillary light reflex was used in Desbeaumes Jodoin et al. (2015) because it provides an opportunity to tease apart sympathetic and parasympathetic influences of iVNS. Similarly, it is of interest to determine whether pupil dilation during taVNS is due to changes in one or both systems. In principle, the use of GAMMs to analyze the data in the present study would allow differences between active and sham taVNS to appear on specific features of the pupillary light reflex and during the prestimulus resting-state period. However, our results indicated that increased pupil diameter during active versus sham taVNS was fairly consistent across the analysis window, and thus do not suggest differential effects of taVNS on sympathetic and parasympathetic activity. While this is largely consistent with the findings reported in Desbeaumes Jodoin et al. (2015) it is possible that differences would be found if analyzing specific features of the pupillary light reflex, such as maximum constriction velocity or maximum constriction acceleration (selectively sensitive to parasympathetic activity), or 75% amplitude recovery time or recovery amplitude after 2.4 sec, which are selectively sensitive to sympathetic activity (see Desbeaumes Jodoin et al., 2015 for details). We did not extract and analyze such features in the present study, because we determined that modeling the entire pupillary response using GAMMs would provide the most informative results for the present aims. Once reliable taVNS parameters are established, it will be more practical to tease apart sympathetic and parasympathetic influences.

It is important to recognize that there is a potential dissociation between the effects of taVNS on pupil size and the desired behavioral outcomes. The link between LC-NE activation via taVNS and target behavioral outcomes, such as improved learning and memory performance, is not direct and finding taVNS parameters that reliably dilate the pupil does not guarantee the same parameters will lead to into improved target outcomes. Animal and human studies involving iVNS have shown that iVNS is most effective at a moderate intensity (∼0.4–0.8 mA), with target behavioral and physiological effects (such as cortical changes) lessening at intensities above and below this range (Borland et al., 2016; Clark et al., 1999). Thus, finding a biomarker for LC-NE activation due to taVNS is a first step. For this biomarker to be useful, further work is needed to establish its relationship to target outcomes. Future work will link the pupil findings in this study to behavioral outcomes.

## 5 CONCLUSION

These present findings contribute to the limited and conflicting evidence that has been found for the relationship between taVNS intensity and pupil dilation. The taVNS parameters used in this study, specifically stimulation of the EAM below perceptual threshold using 50 μs pulses delivered at 300 Hz, have been tested far less than wider taVNS pulses (200–300 μs) delivered at slower rates (25–30 Hz) to other areas of the outer ear, most often the cymba conchae. Across studies, evidence has been mixed regarding the ability of taVNS to effectively engage the LC-NE system, as evidenced by various biomarkers such as pupil size, electrophysiological, and cardiac measures. The results of this study indicate that our stimulation parameters replicate and extend the effects of iVNS in Desbeaumes Jodoin et al. (2015) and the effects of taVNS in Wienke et al. (2023) on overall pupil size during stimulation, reflecting increased LC-NE activity due to taVNS. This finding adds to the evidence suggesting that 300 Hz taVNS applied to the EAM effectively engages the LC-NE system. The present study’s results also suggest a nonlinear relationship between taVNS intensity and resulting increases in pupil size, suggesting that pupillometry may provide a useful physiological biomarker of the effectiveness of taVNS at modulating LC-NE activity. Establishing a reliable biomarker for LC-NE activation via taVNS is needed to titrate stimulation for different applications and individuals and is critical to advancing use of taVNS (Yap et al., 2020).

## 7 AUTHOR NOTES

We thank the study participants. We are grateful to Matthew Turner, Sara McConnell, Aidan Marshall-Cort, and Ben Rickles for their assistance with data collection, Jarrett Lee for assistance programming tVNS, and Henk Haarmann for contributions to earlier portions of the overall project. This material is based upon work supported by the Naval Information Warfare Center and Defense Advanced Research Projects Agency under Cooperative Agreement No. N66001-17-2-4009. For Dr. Phillips and Dr. Kuchinsky: The identification of specific products or scientific instrumentation is considered an integral part of the scientific endeavor and does not constitute endorsement or implied endorsement on the part of the author, DoD, or any component agency. The views expressed in this article are those of the author and do not reflect the official policy of the Department of Army/Navy/Air Force, Department of Defense, or U.S. Government.

## Footnotes

1 Due to equipment malfunction, one participant completed the pupillary light reflex task at the end of the final post-test session of the larger study. Although they received active taVNS during 4 days of training, they had not received taVNS during the 27 days leading up to this session.

2 A firmware update to the DS8R has since allowed intensities lower than 2.0 mA to be reliably delivered.

3 Right-eye pupil dilation in Pandža et al. (2020) was interpreted as an index of attentional effort, rather than LC-NE activity, due to the learning nature of the task.

## Notes

### Competing Interest Statement

The authors have declared no competing interest.

